# Resolving exit strategies of mycobacteria by combining high-pressure freezing with 3D-correlative light and electron microscopy

**DOI:** 10.1101/2023.04.24.538041

**Authors:** Rico Franzkoch, Aby Anand, Leonhard Breitsprecher, Olympia E. Psathaki, Caroline Barisch

## Abstract

The infection course of *Mycobacterium tuberculosis* is highly dynamic and comprises sequential stages that require damaging and crossing of several membranes to enable the translocation of the bacteria into the cytosol or their escape from the host. Many important breakthroughs such as the restriction of vacuolar and cytosolic mycobacteria by the autophagy pathway and the recruitment of sophisticated host repair machineries to the *Mycobacterium*-containing vacuole have been gained in the *Dictyostelium discoideum*/*M. marinum* system. Despite the availability of well-established light and advanced electron microscopy techniques in this system, a correlative approach that integrates both methodologies with almost native ultrastructural preservation is still lacking at the moment. This is most likely due to the low ability of *D. discoideum* to adhere to surfaces, which results in cell loss even after fixation. To address this problem, we improved the adhesion of cells and developed a straightforward and convenient workflow for 3D-correlative light and electron microscopy. This approach includes high-pressure freezing, which is an excellent technique for preserving membranes. Thus, our method allows to monitor the ultrastructural aspects of vacuole escape which is of central importance for the survival and dissemination of bacterial pathogens.

## Introduction

Phagocytosis is a fundamental defence mechanism that triggers inflammatory and immunological responses and plays a crucial role in bridging innate and acquired immunity against invading pathogens (Lim et al., 2017, Lee et al., 2020). Professional phagocytes such as macrophages and neutrophils use a variety of innate immunity strategies to identify and eliminate intruders within bacteria-containing phagosomes. However, some microbes have developed sophisticated strategies to overcome these processes. For example, *Mycobacterium tuberculosis*, the causative agent of tuberculosis, subverts this compartment into a proliferation-friendly niche while also damaging the membrane and rendering the phagosomal defences ineffective. The host, on the other hand, counteracts by recruiting various membrane repair pathways to the site of damage to retain *M. tuberculosis* inside the *Mycobacterium*-containing vacuole (MCV) (Barisch et al., 2023). When the damage overwhelms the repair machineries, *M. tuberculosis* translocates into the cytosol before exiting the cell (Bussi and Gutierrez, 2019).

We and others have pioneered the investigation of various membrane repair pathways such as ESCRT- and autophagy- (López-Jiménez et al., 2018), as well as ER-dependent repair (Anand et al., 2023) in the *Dictyostelium discoideum*/*M. marinum* model. Within this system, the sequential stages of the *M. tuberculosis* infection course such as (i) the initial stage when the bacteria remodel the phagosome, (ii) the vacuolar stage when replication begins, and (iii) vacuole exit and the cytosolic stage that precedes (iv) host cell exit, are conserved and are distinguishable with well-established markers (Figure 1) (Cardenal-Munoz et al., 2017). In *D. discoideum*, light microscopy (LM) of live cells has been increasingly important to study the course of infection due to it’s high time and spatial resolution. For example, LM was used to generate time-lapse movies to monitor the escape of *M. marinum* via ejectosomes (Hagedorn et al., 2009), the spatiotemporal dynamics of Rab proteins at the MCV (Barisch et al., 2015a) and the re-distribution of lipid droplets during infection (Barisch et al., 2015b). Although LM is a powerful tool for observing dynamic processes, it does have limitations. For instance, it relies on selective labelling techniques, e.g. fluorescent proteins or dyes, to visualize subcellular structures or protein localization. Additionally, the relatively low resolution of LM and the absence of ultrastructural context makes it difficult to accurately identify the structures underlying the fluorescence signal. For several decades, transmission electron microscopy (TEM) has been used to unravel the ultrastructural architecture of various organelles. Unlike other techniques, TEM provides near-atomic spatial resolution and enables simultaneous visualization of all subcellular components (Bozzola and Kuo, 2014). In the *D. discoideum*/*M. marinum* system, TEM was used to image ejectosomes (Hagedorn et al., 2009) and to monitor the accumulation of intracytosolic lipid inclusions inside *M. marinum* (Barisch and Soldati, 2017). However, conventional sample preparation for TEM is known to induce artefacts such as cell shrinkage and extraction of cellular material which can drastically alter the ultrastructure of the sample (McDonald and Auer, 2006).

**FIGURE 1.**
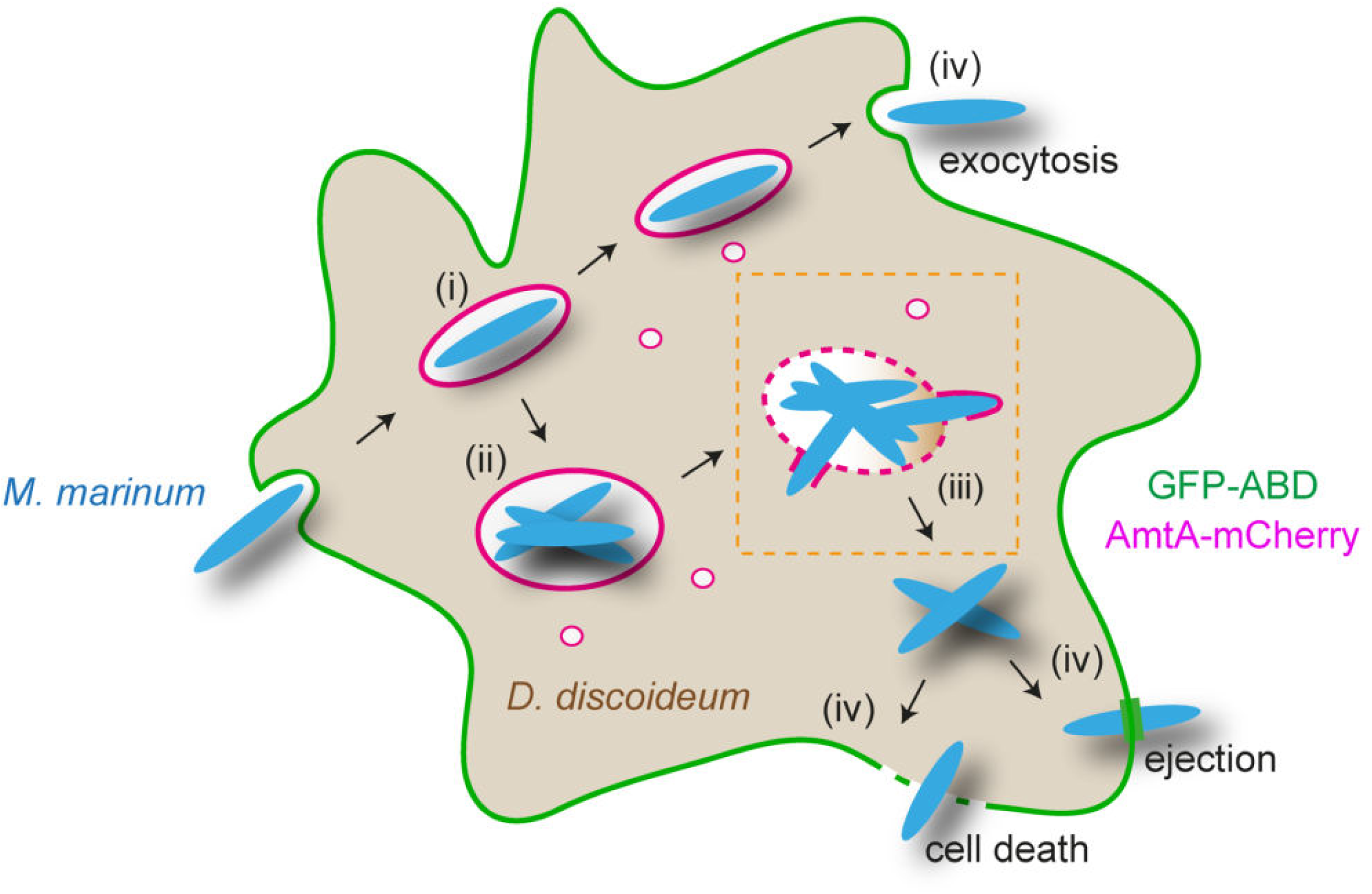
Scheme sketching the sequential infection stages of *M. marinum* in *D. discoideum*. Early after uptake, *M. marinum* resides in the MCV (i) and creates a friendly environment for its own replication by blocking phagosome maturation (ii). Cumulative damage at the MCV membrane leads to vacuole escape of the bacteria (iii). *M. marinum* exits *D. discoideum* either by exocytosis, ejection or upon host cell death (iv). The markers GFP-ABD (F-Actin, cell cortex) and AmtA-mCherry (endosomes and MCV) are suited to distinguish various infection stages. The orange box indicates the “infection stage of interest” that was investigated using the here described workflow.

The combination of both imaging approaches as correlative light and electron microscopy (CLEM) overcomes the individual limitations and thus represents a robust tool that has recently gained increasing attention in the field of infection biology (Kommnick and Hensel, 2021, Weiner et al., 2016b, Lerner et al., 2020). Especially, when investigating vacuolar escape, reliable preservation and visualization of the vacuolar membrane is of utmost importance. To overcome the induction of artefacts during sample preparation, high-pressure freezing (HPF) and freeze substitution (FS) were established already in the 1980s (Moor, 1987, Humbel and Müller, 1985). These techniques reduce the occurrence of artefacts and preserve also membrane organization in a more close to native state (Vanhecke et al., 2008, Kaneko and Walther, 1995). Utilising HPF and FS is therefore essential for accurately detecting vacuolar damage and escaping pathogens. However, an approach that combines HPF and FS with CLEM has yet to be established for the *D. discoideum* system. This is very likely due to the fact that *D. discoideum* adhesion points are small, relatively weak and transient (Mijanović and Weber, 2022). Consequently, the ability of *D. discoideum* to adhere to surfaces is relatively low and is very likely responsible for the great loss of cells during HPF and FS.

The three-dimensional organisation of a cell poses a challenge for the detection of membrane damage through conventional TEM techniques, as they are limited to capturing two-dimensional sections. High-resolution 3D-EM using TEM-tomography has become an essential tool for accurately identifying membrane discontinuity. For instance, in a recent study, membrane rupture within mitochondria during apoptosis was elegantly shown using this approach (Ader et al., 2019). Due to the lack of protocols that allow the visualization of membrane rupture as well as vacuole escape of *M. marinum*, we developed a 3D-HPF/FS-CLEM workflow for the *D. discoideum/M. marinum* system.

To this end, we first improved cell-to-surface adherence of *D. discoideum* before combining high resolution fluorescence microscopy, HPF, and FS to preserve ultrastructure and TEM tomography to get close to atomic resolution. Our protocol thus provides a basis for studying exit events in *D. discoideum*, and other model systems and presents the first published HPF-CLEM protocol for the *D. discoideum/M. marinum* model system.

## Results

### Optimizing D. discoideum adherence and fixation for CLEM

*D. discoideum* adheres to surfaces via focal-like adhesions that resemble the ones from fibroblasts and other mammalian cells. While these points have been shown to be F-actin positive, *D. discoideum* cell-to-surface adhesion includes the homologue of talin A and several integrin-like proteins (Mijanović and Weber, 2022, Kreitmeier et al., 1995, Cornillon et al., 2008). However, adhesion points in *D. discoideum* are characterized by their small size (< 10 % of cell size), fragility, and short lifespan (Wessels et al., 1994, Weber et al., 1995). This has the advantage that passaging *D. discoideum* is relatively easy compared to mammalian cells. Tapping the petri dish or pipetting media onto the cells typically results in cell detachment. Therefore, techniques like immunostaining or EM are challenging as the standard protocols involve many washing steps. To resolve this issue, we set out to optimize the cell-to-surface adherence of *D. discoideum*. We took advantage of the fact that the attachment of cells is improved by enhancing their electrostatic interactions by treating surfaces with poly-l-lysine (PLL) (Mazia et al., 1975). Consequently, cell adherence with and without PLL treatment of sapphire discs, i.e. coverslips suitable for HPF, was evaluated before and after several washing steps using LM. Without treatment, cells detached from large areas of the discs (Supplementary Figure S1), while PLL coating improved cell adherence to a certain extent. It should be noted that PLL treatment has been observed to change the morphology of *D. discoideum*, and thus, we kept incubation times to a minimum of 15 min. However, despite the PLL coating, a high proportion of cells detached after HPF. This was improved when a strain expressing GFP-Actin-binding domain (ABD) was used. These cells are very sticky and hard to detach during cell culture. Currently it is poorly understood, how the expression of GFP-ABD improves cell adhesion, however, this might be linked to the fact that foci enriched in F-actin are the anchorage points of traction forces in *D. discoideum* (Mijanović and Weber, 2022, Iwadate and Yumura, 2008). Thus, overexpression of GFP-ABD might increase adhesion by cross-linking the F-actin cytoskeleton. To rule out that this leads to artefacts, we performed an infection experiment in cells co-expressing AmtA-mCherry (Figure 2). AmtA is an ammonium transporter that labels all the endosomes in *D. discoideum* (Uchikawa et al., 2011) and is a suitable marker for the MCV (Barisch et al., 2015b). While GFP-ABD enabled us to monitor MCVs at early infection stages and host cell exit by ejection (GFP-ABD^+^), AmtA-mCherry allows to visualize MCV membrane rupture and cytosolic bacteria (AmtA-mCherry^−^) (Figure 2). Importantly, the timing of sequential stages of the infection (Figure 1) appeared to be unaltered by the overexpression of both proteins (Figure 2).

**FIGURE 2.**
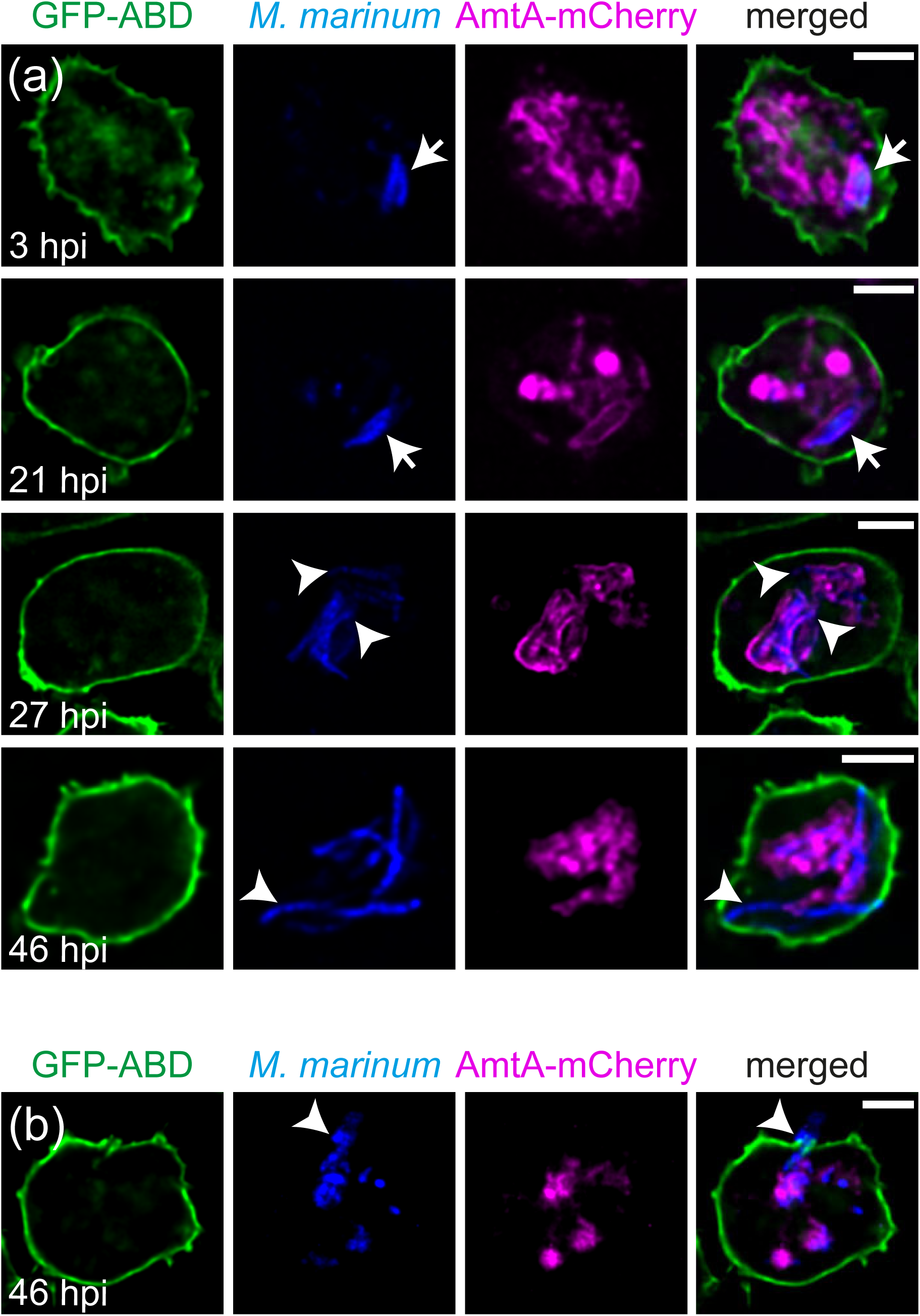
The infection course in cells overexpressing AmtA-mCherry and GFP-ABD. *D. discoideum* overexpressing both, GFP-ABD and AmtA-mcherry, were infected with eBFP-expressing *M. marinum*. At the indicated time points, cells were imaged live. Arrows label vacuolar mycobacteria, arrowheads point to either cytosolic mycobacteria in (a) or a bacterium that is exiting the host by ejection in (b). Scale bars, 5 μm.

In summary, treating sapphires with PLL as well as the overexpression of GFP-ABD significantly improved cell-to-surface adherence of *D. discoideum* and did not impact on the infection course. *D. discoideum* moves a lot and even escapes the imaging field (Barisch et al., 2015a). This, together with the extremely dynamic nature of its organelles including the MCV, and the time lag between LM and HPF, makes the correlation very challenging. To address this, we first tested how different concentrations of the fixative glutaraldehyde (GA) affect cell shape by seeding *D. discoideum* on sapphire discs and transferring them into an ibidi 8-well dish for live imaging. Next, we carefully added double-concentrated fixative to the cells at final concentrations of 0.05 %, 0.1 % and 0.5 % GA and avoided laser exposure to reduce phototoxicity. This revealed that low concentrations of GA eventually immobilized the cells, however, their morphology still continued to change (Supplementary Figure S2a, Movie S1-4). In addition, we often observed cells rounding up especially when incubated with 0.1 and 0.5 % GA. We additionally tried a combination of 0.05 % GA and 3 % paraformaldehyde (PFA) which even lead to the subsequent shrinkage of cells (Movie S5). Finally, we discovered that dipping the sapphire into a solution containing 2% GA for a short exposure, followed by imaging in 0.5% GA, effectively preserved the cell morphology for at least 30 minutes (Supplementary Figure S2a, b, Movie S4). Under these conditions we were able to locate the cells after fixation (Supplementary Figure S2b) and did not observe any interfering background signals (Supplementary Figure S2c). A beneficial side effect of the GA fixation was the even further improved adherence which has already been demonstrated for *D. discoideum* (Koonce et al., 2020).

*Design of a LM configuration that allows the use of high numerical aperture oil immersion objectives* Initially, we imaged upright sapphire discs (3*0.05 mm) with cells on top in an ibidi 3-cm dish or 8-well slide. Using this setup, focusing the cells with inverted 63× or 40 × oil immersion objectives was difficult, and even if it was feasible, the quality of the LM image was poor. We believe that this was caused by (i) the limited working distance of these objectives or by (ii) optical interferences of the bottom of the imaging slide and the sapphire disc.

Subsequently, we designed an imaging setup that is simple to install and could be utilised to acquire LM images of exceptional quality with high NA oil immersion objectives. Figure 3a shows a schematic representation of the entire setup. First, a coordinate system was sputtered onto acid cleaned sapphire discs (3*0.16 mm) using a 20 nm thin layer of gold. Following that, a gold grid with a 2 mm aperture was attached on top of the sapphire disc using Loctite AA 350 to avoid crushing of the cells during the imaging process (Figure 3b). This glue has proven to be non-toxic to cells, is easily applicable and non-soluble in water or acetone (Brown et al., 2012). The assembly was UV-cured overnight in a Leica AFS2 and then coated with PLL. Cells were seeded on top of the sapphire disc with gold spacer at appropriate densities. We used extra thin 24-mm coverslips (130 — 160 μm) as a bottom for the construct, which was fitted into a custom-made holder onto the microscope stage and sealed with a rubber ring (Figure 3c). To prevent cells from drying, 500 μl of Hl5c filtered medium containing 0.5 % GA were added. The sapphire disc was then flipped upside down and placed on the 24 mm coverslip, resulting in cells directly facing the objective (Figure 3a). This enabled high resolution LM imaging using the 63x 1.46 NA oil immersion lens.

**FIGURE 3.**
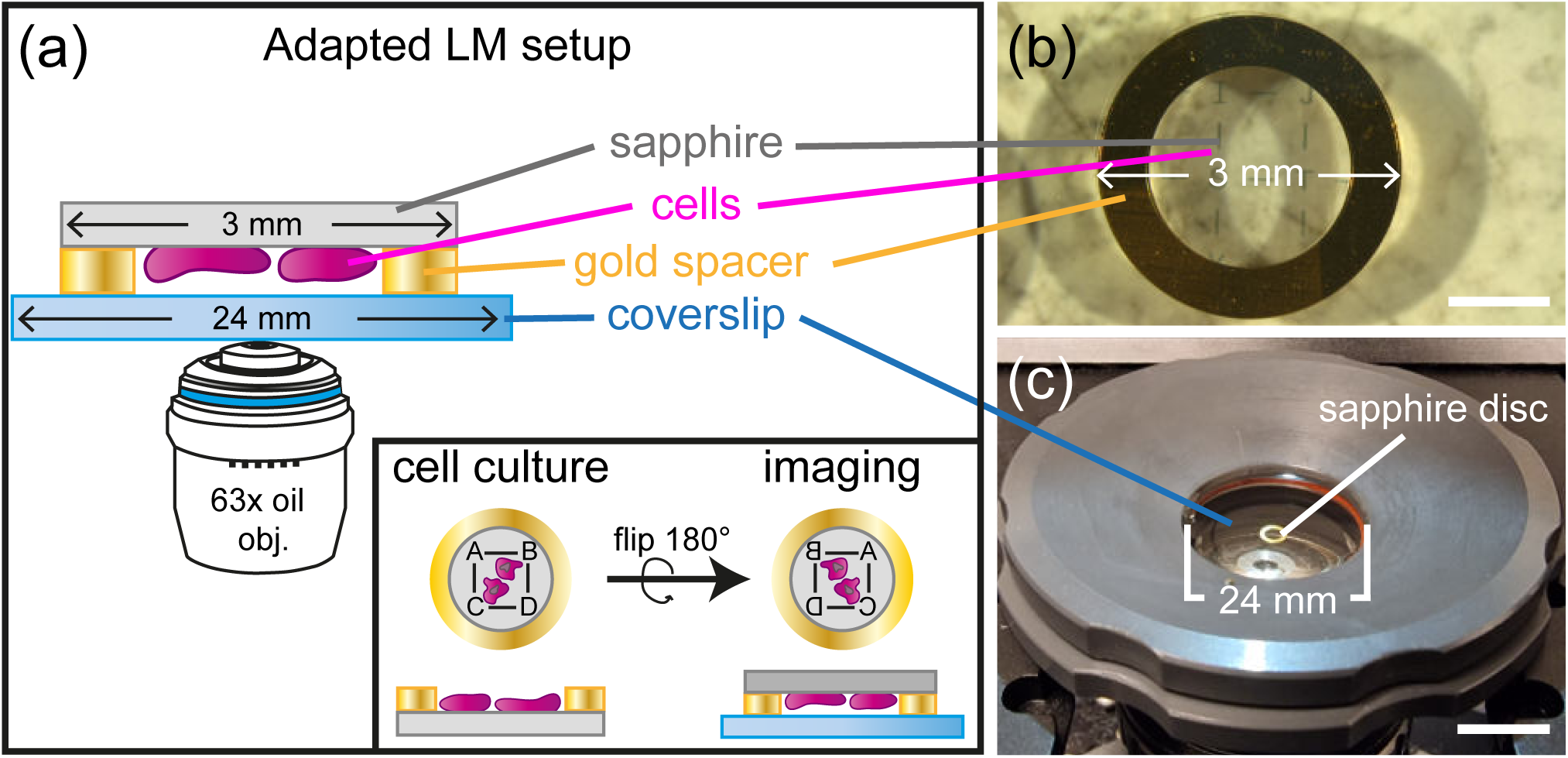
Light microscopy imaging setup for optimized z-resolution and correlative work flows. (a) Schematic representation of a 3 mm sapphire disc (grey) with gold spacer (yellow) and monolayer of cells (magenta) on a 24 mm coverslip facing the 63× NA 1.46 oil immersion objective. Rotation of the disc (as indicated in the inset) and application of the gold spacer allows reliable and reproducible focusing since the laser has to pass only through the coverslip. (b) Customization of the sapphire discs. Cells are seeded on sapphire discs with coordinate system until they form a monolayer of about 70% confluency. (c) Position of the sapphire discs on the stage. The sapphire discs are flipped and placed on a coverslip that is mounted with the help of a custom-made adaptor.

### Workflow of 3D-HPF/FS-CLEM of infected D. discoideum

The implementation of our improved cell adhesion and imaging protocols, combined with HPF and FS resulted in highly reproducible outcomes. The whole procedure is described in the following: Briefly, infected cells expressing ABD-GFP and AmtA-mCherry were seeded on PLL-coated sapphire discs. Prior to LM, the discs were dipped into 2% GA (in Hl5c filtered medium) and transferred upside down with the cells facing the coverslip into the custom-made coverslip holder (Figure 3). Samples were screened for events of interest and a high-resolution z-stack (Figure 4a) as well as a low resolution brightfield overview were acquired (Figure 4b). Following this protocol, at least two positions were monitored on each sapphire disc. Directly after acquisition of the last image, the sapphires were transferred into the HPF holder with the cells facing upwards (Figure 4c, i). The flat side of a planchette was dipped into 1-hexadecene and added on top. The assembly was closed and subjected to HPF (Figure 4c, ii). After HPF, the samples were stored in specially designed vessels in liquid nitrogen. These vessels consisted of a 0.5 ml tube with cut-off top and bottom and were used from this point on during the whole FS and embedding procedure. To prevent samples from slipping through the bottom, while also allowing solutions to access the specimen, a fine piece of mesh was attached by slightly melting the end of the tube and pressing it onto the mesh (Figure 4d). This avoids problems that occur when samples are moved to fresh tubes for each step and turned out to be critical in preventing cells from detaching from the sapphires. For FS we chose a protocol that we established previously and worked well for a range of different samples in our facility (see Experimental Procedures). For final polymerization in EPON 812, sapphire discs were transferred with the cells facing upwards into a 0.2 ml tube which was then filled with resin (Figure 4e, i). Polymerized blocks were separated from the tubes using a razor blade and excess resin on the top and sides of the sapphire was removed. By immersing the top of the block into liquid nitrogen and then touching the sapphire disc with a 50-degree warm razor blade, the disc was removed and the EPON surface with cells and coordinate system became visible (Figure 4e, ii). To facilitate trimming and initial correlation, images of the whole block-face were acquired in a scanning EM (SEM) using a low vacuum mode with high probe current and voltage (Figure 4f, g). This revealed the gold imprint of the coordinate system (Figure 4g) and confirmed that the cells remained on the sapphire during the whole procedure thus making a first correlation possible (Figure 4h-j). In addition, this allowed precise targeting and trimming of the block to exclusively include the region of interest (ROI) (Figure 4k). 250-nm-thin sections were cut using an ultramicrotome and deposited on copper slot grids (Figure 4l). With the help of LM and SEM data, the cell of interest was rapidly identified and relocated on the section (Figure 4k). After addition of gold fiducials and contrasting with lead citrate and uranyl acetate correlative TEM tomograms were acquired (Figure 4l).

**FIGURE 4.**
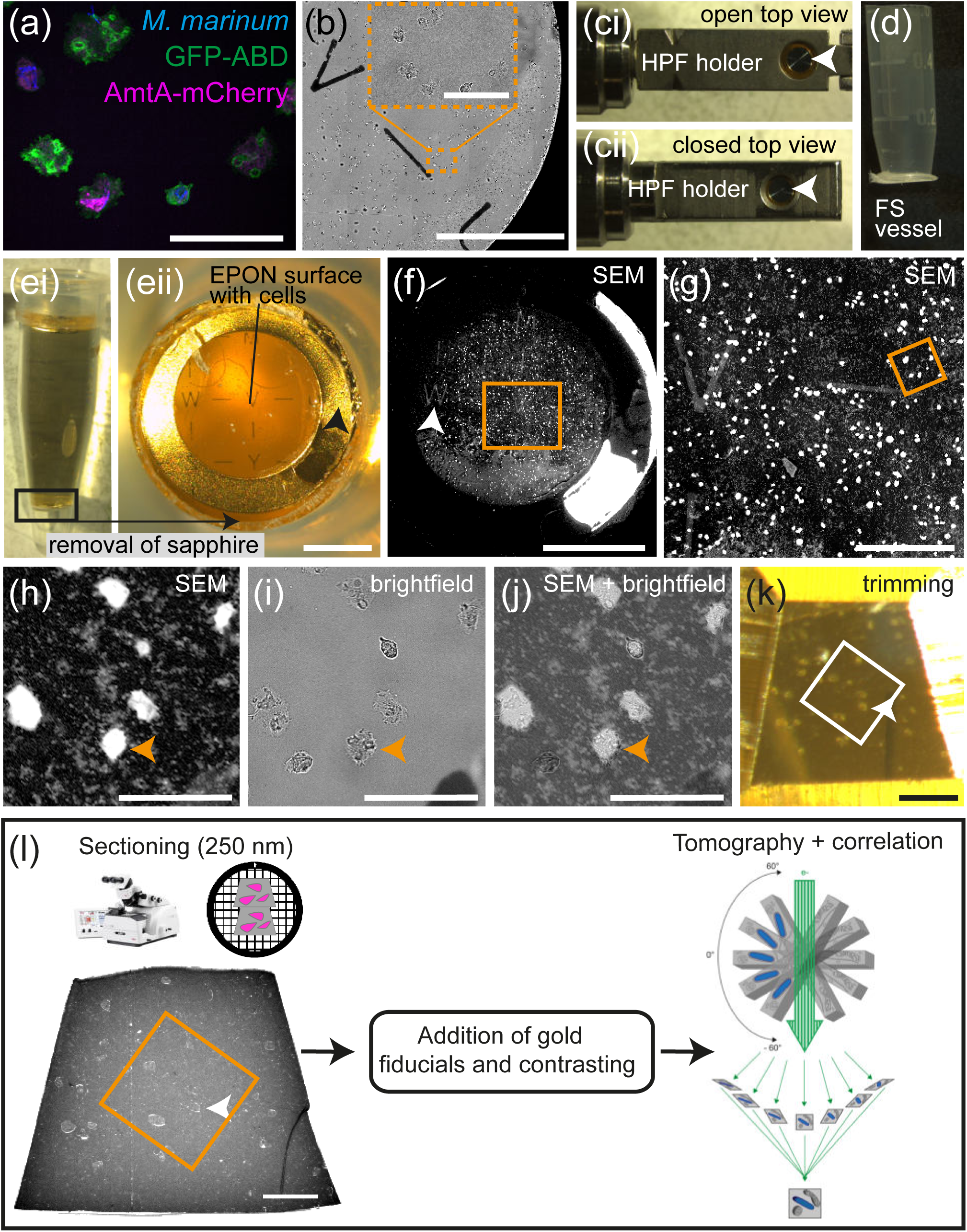
3D-HPF/FS-CLEM workflow. (a) Locating the cell of interest and fluorescence z-stack acquisition of fixed cells by SD microscopy using the 63× objective. (b) An overview is acquired using either the 25× oil immersion or 10x air objective to visualize the coordinates of the desired region (orange box). (c) Assembly of the sandwich consisting of a flat planchette dipped in 1-hexadecene and the sapphire with gold spacer and cells is assembled in the HPF-holder (i). With the redesigned carrier, the holder can be closed properly and HPF can be initiated (ii). (d) After HPF, sapphire discs are placed in a special vessel that facilitates storage in liquid nitrogen and their further processing. (e) Embedding of cells in a 0.2 ml tube that is filled with EPON 812 (i) and removal of the sapphire via heat shock which leaves the resin with cells, the gold spacer and the coordinates on top (ii, black arrowhead). (eii) Bottoms-up view of (ei). (f-j) Locating the cells of interest (orange arrowheads) by acquiring an overview using SEM in which the gold coordinates are easily visible (f, white arrowhead), zoom of orange box in f (g), zoom of orange box in g (h), bright field image of the SEM region shown in h (i) and its overlay (j). (k) Trimming of the ROI and sectioning of 250 nm thin sections using an ultramicrotome facilitated by the previously recorded LM and SEM images. Sections are then transferred onto copper grids. (l) Addition of gold fiducials and contrasting agents prior to tomogram acquisition and correlation. Arrowheads in (eii), (f), (h) - (l) indicate the cell of interest. White or orange boxes in (k) and (l) indicate the ROI. Scale bar, 50 μ m in (a), (h), (i), (j); 500 μm in (b); 1 mm in (e; i, ii); 200 μm in (g) and (l); 100 μm in (k).

In conclusion, our approach enables the efficient correlation of LM and EM data, and should consequently facilitate an accurate and comprehensive understanding of subcellular structures and dynamic processes including vacuole escape and host cell exit.

*3D-HPF/FS-CLEM to monitor the ultrastructure of vacuolar escape of M. marinum in D. discoideum* To showcase the applicability of our developed 3D-HPF/FS-CLEM workflow, we set out to resolve and correlate the ultrastructure of the MCV during vacuolar escape of *M. marinum* in *D. discoideum*. Cells co-expressing GFP-ABD and AmtA-mCherry were infected with eBFP-expressing bacteria and analysed at 24 hours post-infection (hpi) (Figure 5). During LM we partly observed absence of the AmtA-mCherry signal on the MCV indicating that there might be membrane rupture at the site of escape (Figure 5a, arrowheads). However, without the underlying ultrastructural information, this interpretation remains speculative. With the help of the previously described 3D-HPF/FS-CLEM workflow, we were able to precisely correlate areas without AmtA-mCherry signal with high resolution tomography data (Figure 5b, Movie S6-7). This revealed one single MCV containing at the same time a large membrane rupture with partially cytosolic bacteria (Figure 5c), as well as a small membrane lesion in close proximity to vacuolar bacteria (Figure 5d). Thus, the developed 3D-HPF/FS-CLEM workflow drastically facilitated monitoring of membrane discontinuities at the MCV without the risk of interpreting artefacts. In conclusion, our protocol allows the visualisation of the ultrastructure of vacuolar exit sites of mycobacteria and very likely also other pathogens and may be in addition able to resolve bacteria uptake as well as host cell escape in the future.

**FIGURE 5.**
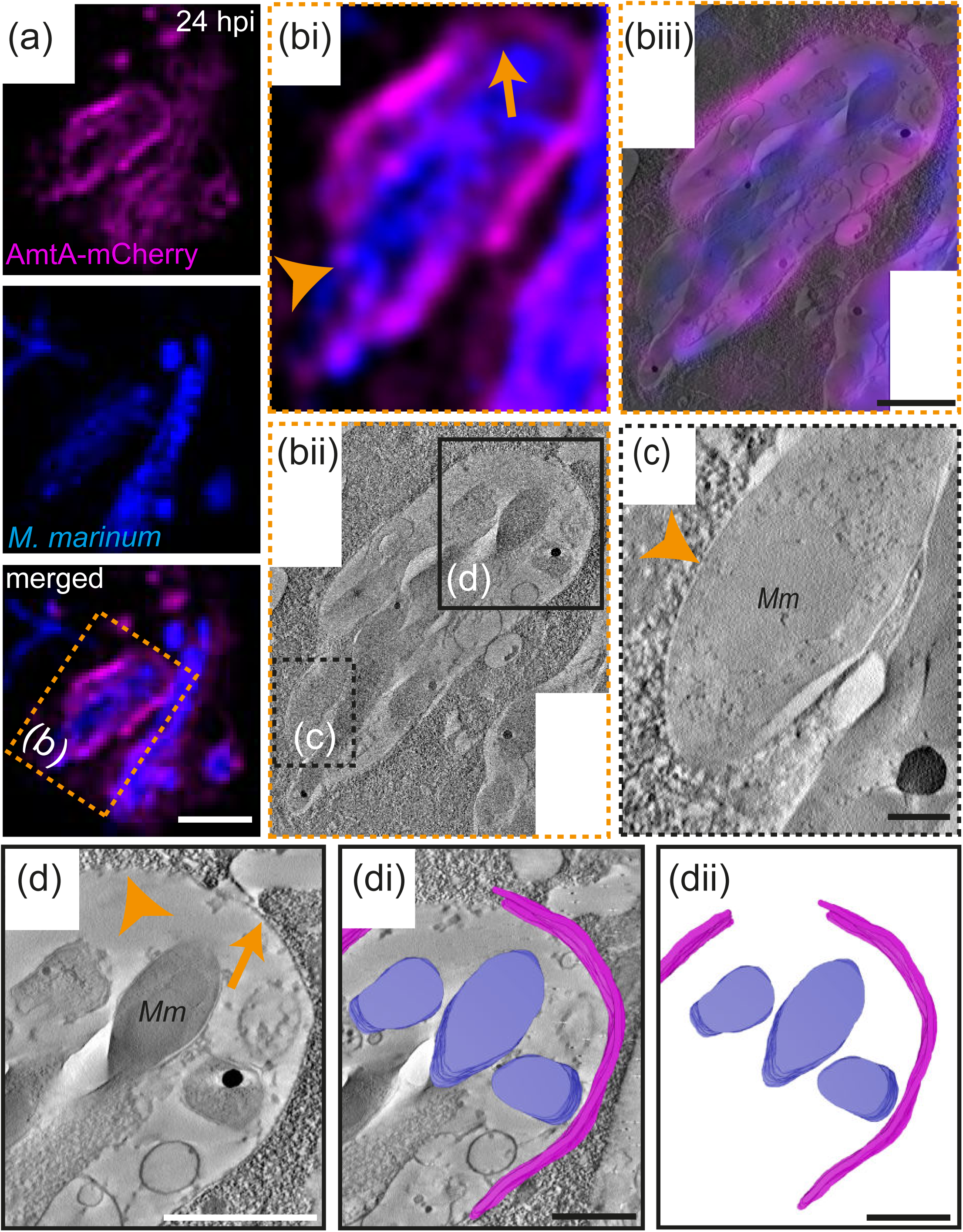
3D-HPF/FS-CLEM enables the visualization of *M. marinum* exiting the MCV. (a) Fluorescence image showing the cytosolic translocation of *M. marinum*. (b) Correlation of the magnified LM image with the tomogram; (i) LM image showing membrane ruptures reflected by the loss of the AmtA-mCherry-signal (arrow: minor lesion, arrowhead: full rupture). (ii) Corresponding tomography slice. (iii) Overlay of both imaging modalities to identify possible exit events. (c-d) Higher magnification tomograms of the areas indicated in (bii) revealed bacterium that is fully cytosolic (c, arrowhead) and bacteria that are partially enclosed by the membrane of the MCV (arrow). (d) The gap in the MCV membrane correlates with the absence of the AmtA-mCherry-signal (arrowhead). (di, dii) This is supported by the segmentation showing a discontinuity of the MCV membrane throughout the whole tomogram. Blue: *M. marinum*, magenta: MCV membrane. *Mm*: *M. marinum*. Scale bar, 5 μm (a), 1 μm (b), 200 nm (c), 1 μm (d). Please see also Movie S6-S7.

## Discussion

In this article we describe a robust 3D-HPF/FS-CLEM workflow that integrates LM, HPF, FS and TEM-tomography for studying exit events in the *D. discoideum*/*M. marinum* model system. To this end, we created a unique LM setup which facilitates high resolution LM using high numerical aperture oil immersion lenses that is also compatible with live cell imaging (Figure 3, Supplementary Movies S1-5). However, because of the time delay of up to 1 min between acquisition of the last LM image and HPF, we decided to include a mild fixation step. This immobilizes *D. discoideum* and allows precise correlation of the LM and the TEM-tomogram. Alternatively, the high-pressure freezer could be placed directly adjacent to the LM. Moreover, to enable fast and easy transfer of the sample, a system was developed, in which LM and HPF are placed in the same holder. Using this system, time until cryo-fixation can be reduced to four seconds (Brown et al., 2012, Verkade, 2008). This configuration is only available from Leica which severely limits its use. Another comparable setup was recently presented by Heiligenstein *et al*. consisting of the HPM live μ high-pressure freezer in combination with the CryoCapsule (Heiligenstein et al., 2021). Here, the process from LM to cryo-fixation is completely automated and happens within 1.2 seconds. These developments underline the urgent need for new methodological approaches for sample carriers and preparation for CLEM including HPF. In this regard we developed a system which is easy and cheap to setup, does not require special equipment and allows precise correlation of events monitored by LM (Figure 3-4). However, there is still room for improvement with regard to the quick transfer between LM and HPF. Additional modifications of our method would include the design of an LM setup in which LM is directly carried out through the bottom of the sapphire rather than the coverslip. Because the cells would be closer to the objective lens, higher-quality fluorescence images would most likely be obtained, but construction and handling of such a system would be even more challenging.

Key advantages of utilizing HPF and FS include the reduction of artefacts and also the overall superior preservation of ultrastructure including membranes which present important barriers pathogens have to overcome (McDonald and Auer, 2006, Vanhecke et al., 2008, Kaneko and Walther, 1995). For the *D. discoideum/M. marinum* model system only few CLEM approaches are published (Gerstenmaier et al., 2015, Malinovska et al., 2015). They all include conventional sample preparation for EM which may not accurately reflect the native membrane organization in the investigated specimens and complicates data interpretation. A general challenge for setting up CLEM with *D. discoideum* is the fact that these cells adhere only weakly to the substrate (Wessels et al., 1994, Weber et al., 1995). To overcome this, we used (i) a strain expressing GPF-ABD in which cell-to-substrate adherence is increased (ii) in combination with PLL-coating of the sapphire discs and (iii) a mild GA fixation which drastically improved the success rate of the protocol. Coating substrates with PLL is a well-known method to improve adherence in cell culture and also provided excellent results for *D. discoideum* (Lombardi et al., 2008). Fixation with low concentrations of GA has been previously shown to improve cell attachment in *D. discoideum* and was suggested to be used when several washing or incubation steps are involved (Koonce et al., 2020). Even when chemical fixation is employed prior to HPF, ultrastructural preservation is drastically enhanced, indicating that the most problematic steps in the conventional sample preparation protocol are indeed post-fixation and especially dehydration. This is applicable for a wide range of specimens ranging from highly labile tissues to cells infected with Hepatitis C virus (Romero-Brey and Bartenschlager, 2015, Sosinsky et al., 2008).

Since TEM-tomography is able to achieve high resolution 3D-data for relatively small volumes, a great improvement to understand escape of pathogens would be to also visualize whole cells or larger volumes. For conventional sample preparation this has been already shown for *M. marinum* infected *D. discoideum* and other host-pathogen systems (Weiner et al., 2016a, López-Jiménez et al., 2018, Anand et al., 2023). In principle, specimens prepared with the workflow developed in this study could also be directly processed via focused ion beam (FIB)-SEM to provide high volumetric information with EM resolution (Villinger et al., 2012). This further underlines the versatility of our approach and would allow also correlation in three dimensions. Because EM allows us to see not just one but all cellular compartments, we can conduct correlative studies on other organelles that potentially interact directly with pathogens or the MCV, such as for example the endoplasmic reticulum (Anand et al., 2023).

Altogether, our workflow provides the first approach of correlatively visualizing adherent *D. discoideum* cells infected with *M. marinum* in a more native state utilizing HPF and FS. This makes it possible to evaluate data more accurately and to detect vacuole escape with near-atomic spatial resolution in three dimensions. Other major advantages of our approach are that it is inexpensive and relatively simple to implement and might thus help to monitor membrane damage and exit events in other infection systems.

### Experimental Procedures

#### D. discoideum strains and cell culture

All the *D. discoideum* material is listed in Table 1. *D. discoideum* wild type (AX2) was cultured axenically at 22 °C in Hl5c medium (Foremedium) in the presence of 100 U/mL penicillin and 100 μg/mL streptomycin, respectively. The GFP-ABD expressing cells were electroporated with pDM1044-AmtA-mCherry (Barisch et al., 2015b) and grown in Hl5c containing the respective antibiotics (hygromycin 50μg/ml and G418 at 5 μg/ml).

**Table 1.** Material used in this publication.

| *D. discoideum* strains | Relevant characteristics | Source/Reference |
| --- | --- | --- |
| AX2 | wild type | www.dictybase.org |
| *D. discoideum* Plasmids | Relevant characteristics | Source/Reference |
| GFP-ABD | pDXA-GFP-ABD120, G418<sup>r</sup>, Amp<sup>r</sup> | (Pang et al., 1998) |
| AmtA-mCherry | pDM1044-AmtA-mCherry, Hyg<sup>r</sup>, Amp<sup>r</sup> |  |
| *M. marinum* strains | Relevant characteristics | Source/Reference |
| *M. marinum* M | wt, parental strain | L. Ramakrishnan<br>(University of Cambridge) |
| Mycobacteria Plasmids | Relevant characteristics | Source/Reference |
| pTEC18 | eBFP2 under control of the MSP promoter, Hyg<sup>r</sup>, Amp<sup>r</sup> | Addgene #30177<br>(Takaki et al., 2013) |

#### Mycobacteria strains and cell culture

*M. marinum* was cultured in 7H9 medium supplemented with 10 % OADC, 0.2 % glycerol and 0.05 % Tween-80 at 32 °C in shaking at 150 rpm until an OD600 of 1 (∼1.5 × 10^8^ bacteria/ml). To prevent bacteria from clumping, flasks containing 5 mm glass beads were used. To generate wt mycobacteria expressing eBFP, the unlabelled strain was transformed with the pTEC18 plasmid and cultured in medium containing 25 μg/ml hygromycin.

#### D. discoideum infection with M. marinum

To prepare *D. discoideum* for infection, cells were grown overnight in 10-cm dishes in media devoid of antibiotics until confluency. The infection was carried out as previously described (Hagedorn and Soldati, 2007, Barisch et al., 2015a). Briefly, for a final multiplicity of infection (MOI) of 10, 5 × 10^8^ bacteria were washed twice and resuspended in 500 μl Hl5c. To remove clumps, bacteria were passed 10 times through a 25-gauge needle and added to a 10-cm dish of *D. discoideum* cells. To increase the phagocytosis efficiency, the plates were centrifuged at 500 g for two times for 10 min at RT. After 20–30 min, the extracellular bacteria were removed by several washes with Hl5c. Finally, the infected cells were taken up in 30 ml of Hl5c at a density of 1 × 10^6^ cells/ml supplemented with 5 μg/ml streptomycin and 5 U/ml penicillin to prevent growth of extracellular bacteria and incubated at 25 °C at 130 rpm. At 24 hpi, samples were taken and prepared for live imaging.

#### Preparation of sapphire discs

Before usage, sapphire discs (3mm × 0.16 mm) were acid cleaned. To this end, the discs were soaked in 3.7 % hydrochloric acid (HCl) for an hour on a tumbler, washed with de-ionised water, with 100 % ethanol and air dried. Next, sapphires were equipped with a coordinate system by gold sputtering and a gold spacer to optimize image acquisition. The spacer consisted of a 3-mm gold grid (G2620A, Plano) with a central aperture of 2 mm and was glued onto the sapphire disc using Loctite AA 350 (Henkel, Hemel Hempstead, UK). The glue was polymerized overnight in a Leica AFS2. To improve cell adherence, sapphires were then incubated with 1 mg/ml PLL (P1274, Sigma) for 3 hrs at 37 °C. Unbound PLL was removed by three washes with double distilled water, air dried on Kimtech paper (#7552, Diagonal) and stored at RT.

#### Pre-fixation and fluorescence imaging by spinning disc confocal microscopy

Before seeding infected cells, 6 –8 sapphire discs were washed twice with Hl5c (without antibiotics), placed in a 2-well μ-ibidi dish and checked under a light microscope for handling errors (flipped upside down?). 70 % confluent sapphires were prepared by adding the cells and frequent inspection with a cell culture microscope. To remove floating cells, the medium was then replaced with fresh Hl5c filtered medium. Prior to image acquisition, the sapphires with cells were briefly dipped into 2% GA in Hl5c filtered and transferred with the cells and gold spacer facing the objective into a custom holder containing 0.5 % GA in Hl5c filtered for light microscopy.

The Zeiss Cell Observer.Z1 is an inverted microscope, fully motorized and equipped with a Yokogawa Spinning Disc Unit CSU-X1a 5000 with a custom-built acrylic glass incubation chamber. The chamber was set to 25 °C 30 min prior to image acquisition. Alpha Plan-Apochromat 63x (NA 1.46, TIRF, oil immersion equipped with DIC slider EC PN 63x 1.25 III, CA 63x /1.2 W III), Plan-Apochromat 40× (NA 1.4, DIC, oil immersion equipped with DIC slider CA 40× /1.2 W, LD CA 40× /1.1 W III), LD LCI Plan-Apochromat 25× (NA 0.8, DIC, immersion: oil, water, and glycerin, equipped with DIC slider LCI PN 25×/0.8 II) and Plan-Neofluar 10× (NA. 0.3, DIC I, Ph 1, air) were used as objectives. The system is equipped with computer-controlled multi-color laser module with AOTF combiner (405 nm diode laser, max. power 50 mW, 488 nm optically pumped semiconductor laser, max. power 100 mW, 561 nm diode laser, max. power 40 mW, 635 nm diode laser, max. power 30 mW). The photometrics Evolve EMCCD camera was used and contained filters for blue (Zeiss Filter set 49), eGFP (Zeiss Filter set 38 HE) and Cy3 (Zeiss Filter set 43 HE). An area of 512 x 512 pixels with binning 1×1 was imaged with laser power of 20% to 30% for 488 nm, 10% to 20% for 405 nm and 40% to 50% for 561 nm. All channels were captured with a 100 ms exposure time. Data was acquired in Zeiss ZEN 2012 (blue edition) and stored in the OMERO 5.6.4 database. Images were processed using FIJI/ImageJ and deconvolved using the Huygens Software from Scientific Volume Imaging (Netherlands).

#### High pressure freezing (HPF)

Directly after acquisition of the last light microscopy image, sapphire discs were further processed for high-pressure freezing (HPF). To this end, the discs were placed with the cells and gold spacer facing the flat side of a 3 mm aluminum planchette no. 242 (Engineering Office M. Wohlwend GmbH, Sennwald, Switzerland) which was dipped into 1-hexadecene (822064, Merck). Subsequently, the assembly was placed in the HPF-holder and immediately frozen using a Wohlwend HPF Compact 03 high-pressure freezer (Engineering Office M. Wohlwend GmbH, Sennwald, Switzerland). The vitrified samples were stored in homemade storage vessels consisting of 0.5 ml tubes with cut-off tops and bottoms. A fine mesh was bonded to the bottom to allow the flow of solutions to the samples while preventing them from falling.

#### Freeze substitution (FS)

Prior to FS, the aluminum planchettes were separated from the sapphire discs in liquid nitrogen. All steps were conducted with sapphire discs in aforementioned vessels. The sapphire discs were then immersed in a substitution solution containing 1 % osmium tetroxide (19134, Electron Microscopy Sciences), 0.1 % uranyl acetate (E22400, Electron Microscopy Sciences) and 5 % H2O in anhydrous acetone (83683.230, VWR) pre-cooled to −90 °C. The FS was performed in a Leica AFS2 (Leica, Wetzlar, Germany): 27 hrs at −90 °C, 12 hrs at −60 °C, 12 hrs at −30 °C and 1 hr at 0 °C. After 5 washes with anhydrous acetone on ice, the discs were stepwise embedded in EPON 812 (Roth, Karlsruhe, Germany), incubated with acetone (30% EPON, 60% EPON, 100% EPON) and finally polymerized for 48 hrs at 60 °C. Semithin sections of 250 nm were cut with a Leica UC7 ultramicrotome using diamond knives (Diatome, Switzerland). Sections were collected on formvar-coated grids and post-stained for 30 min with 2% uranyl acetate and 20 min in 3 % lead citrate and analyzed with a JEM 2100-Plus (Jeol, Japan) operating at 200 kV equipped with a 20-megapixel CMOS XAROSA camera (EMSIS, Muenster, Germany).

#### Tomography acquisition

For TEM tomography, 250 nm thick sections were labelled with 10 or 15 nm protein-A-gold fiducials on both sides prior to post-contrasting. Tilt series were acquired from +60° to −60° with 1 ° increments using the TEMography software (Jeol, Japan) and a JEM 2100-Plus operating at 200 kV and equipped with a 20-megapixel CMOS XAROSA camera (EMSIS, Muenster, Germany). Nominal magnifications and pixel size were 10000 x and 0.94 nm (Figure 5b-e) and 15000 x and 0.62 nm (Figure 5f).Tomograms were reconstructed using the back projection algorithm in IMOD (Kremer et al., 1996). Segmentation was done manually using IMOD or Microscopy Image Browser (Belevich et al., 2016) and animations were visualized using Amira-Avizo (Thermo Fisher Scientific, Waltham, USA). For visualisation of the MCV in Figure 5 b-e two independent tomograms were stitched together using Amira-Avizo.

## Acknowledgments

We greatly acknowledge the light microscopy unit of the integrated Bioimaging facility (iBiOs) at the University of Osnabrück and especially Rainer Kurre and Michael Holtmannspötter for their expertise and friendly support. This project was supported by the DFG: SPP2225 (CB: BA 6734/2-1), SFB944 (CB: P25) and SFB1557 (CB: P1). We especially thank Michael Hensel for supporting this work via the iBiOs imaging platform that is part of the SPP2225 (MH: HE 1964/24-1).

## Author Contributions

Rico Franzkoch: Conceptualization; Investigation; Writing - original draft; Methodology; Validation; Visualization; Writing - review & editing; Formal analysis; Software; Data curation.

Aby Anand: Conceptualization; Investigation; Writing - original draft; Methodology; Validation; Visualization; Writing - review & editing; Formal analysis; Software; Data curation

Leonhard Breitsprecher: Visualization; Data curation; Software

Olympia E. Psathaki: Conceptualization; Funding acquisition; Validation; Writing - review & editing; Project administration; Supervision; Resources

Caroline Barisch: Conceptualization; Funding acquisition; Writing - original draft; Validation; Writing - review & editing; Project administration; Supervision; Resources

## Supplementary Material

**FIGURE S1.**
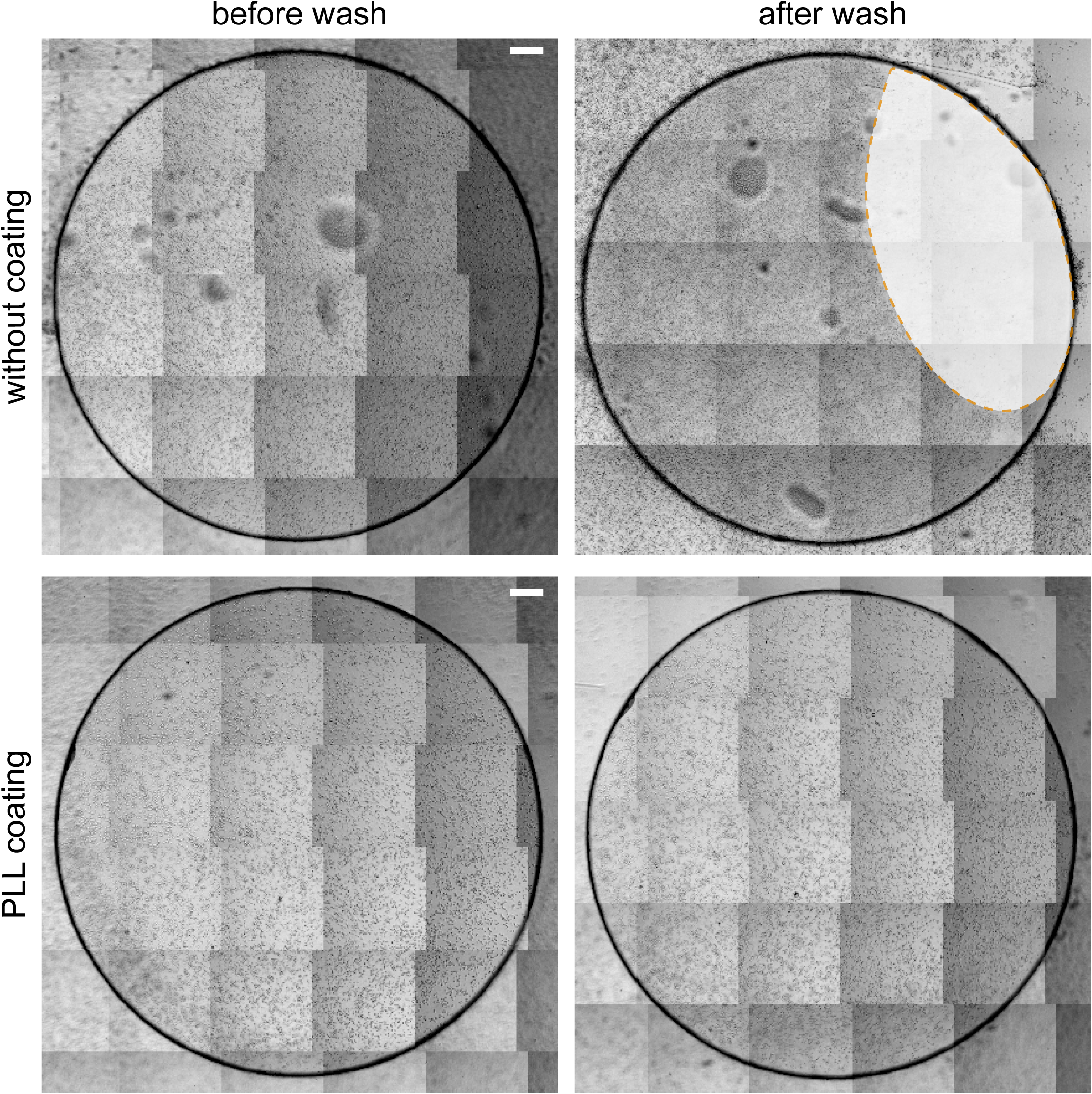
PLL improves *D. discoideum* adherence on sapphire discs. *D. discoideum* wild type was seeded on non-coated or PLL-coated sapphire discs and imaged live before and after washing with Hl5c. An area in which many cells were lost is indicated (orange line). Scale bar, 200 μm.

**FIGURE S2.**
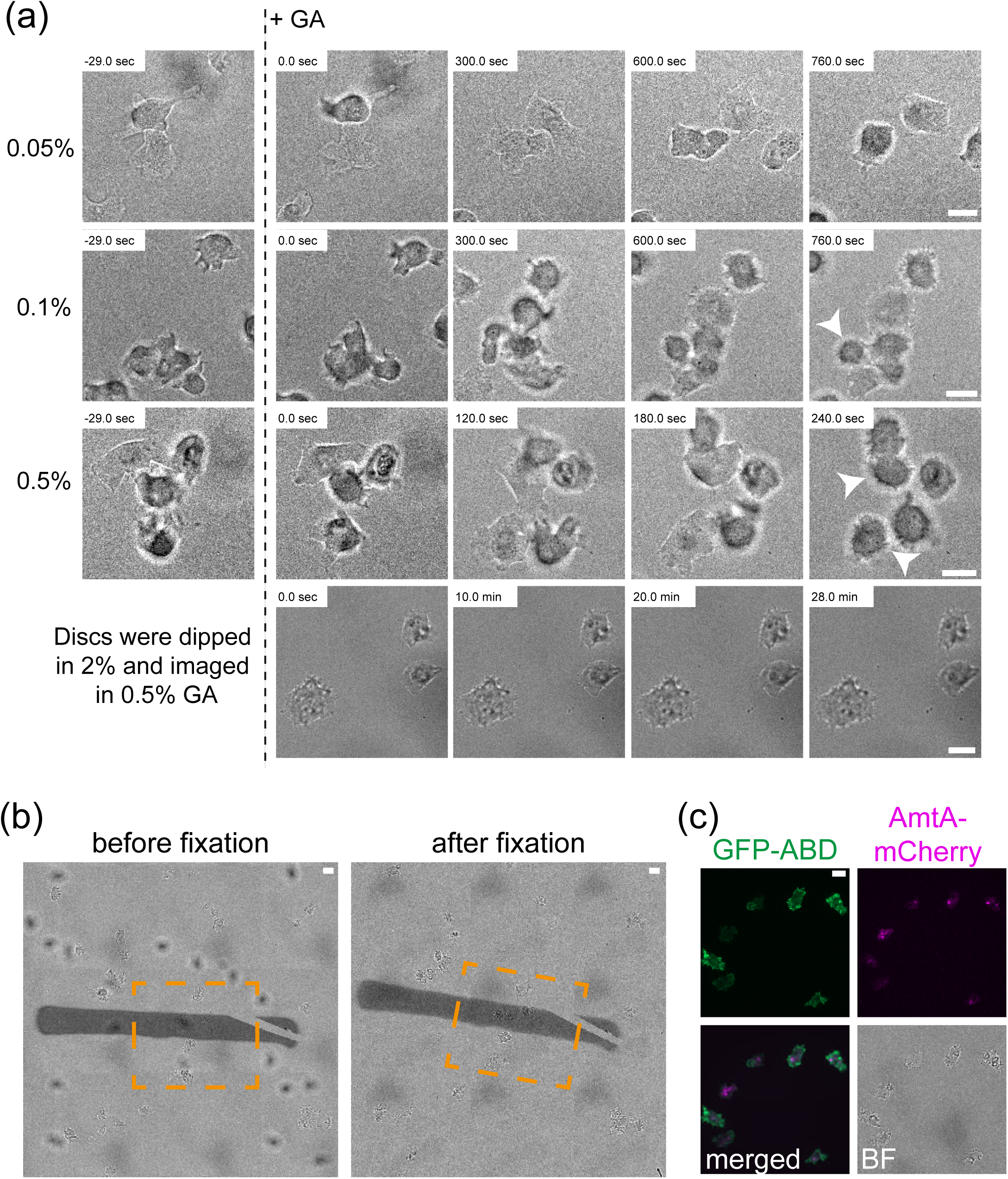
Pre-chemical fixation immobilizes *D. discoideum* and does not impact on background fluorescence. (a) Cell fixation upon exposure to 0.05 %, 0.1 %, 0.5 % GA or upon short immersion in 2 % GA. Incubation with 0.1 % or 0.5 % GA leads to cell death (round cells) while a short immersion in 2 % GA immobilized the cells up to 30 min in 0.5 % GA. (b) Relocation and fluorescence imaging of cells after fixation. (c) 0.5 % GA does not produce any background noise. Cells expressing GFP-ABD and AmtA-mCherry were seeded on sapphires with or without coordinates before movies were recorded. Arrowheads indicate round cells. After fixation, cells can be tracked back (orange box). Fixed GFP-ABD cells were imaged live using the SD microscope with the 63x objective. Please see also Movies S1-5. Scale bar, 10 μm.

**MOVIES S1-S5** Time lapse movies of *D. discoideum* in presence of 0.05, 0.1, 0.5,2 % GA and 3 % PFA and 0.05 % GA. For more information see Figure S2.

**MOVIES S6-S7** Tomogram of the ruptured MCV containing several bacteria as shown in Figure 5. For more information see Figure 5.

